# A Bayesian nonparametric approach to super-resolution single-molecule localization

**DOI:** 10.1101/2020.02.15.950873

**Authors:** Mariano I. Gabitto, Herve Marie-Nelly, Ari Pakman, Andras Pataki, Xavier Darzacq, Michael I. Jordan

## Abstract

We consider the problem of single-molecule identification in super-resolution microscopy. Super-resolution microscopy overcomes the diffraction limit by localizing individual fluorescing molecules in a field of view. This is particularly difficult since each individual molecule appears and disappears randomly across time and because the total number of molecules in the field of view is unknown. Additionally, data sets acquired with super-resolution microscopes can contain a large number of spurious fluorescent fluctuations caused by background noise.

To address these problems, we present a Bayesian nonparametric framework capable of identifying individual emitting molecules in super-resolved time series. We tackle the localization problem in the case in which each individual molecule is already localized in space. First, we collapse observations in time and develop a fast algorithm that builds upon the Dirichlet process. Next, we augment the model to account for the temporal aspect of fluorophore photo-physics. Finally, we assess the performance of our methods with ground-truth data sets having known biological structure.

## 1. Introduction

Super-resolution microscopy (SRM) is an imaging methodology that allows researchers to overcome the diffraction limit imposed by conventional light microscopy [Betzig et al., 2006; Rust, Bates and Zhuang, 2006]. SRM resolves photoswitchable fluorophores in a field of view by sparsely and randomly activating individual emitters and then localizing them with sub-diffraction precision. This technique has the potential to revolutionize the field of cellular microscopy by enabling the study of intracellular proteins within cellular compartments at nanometer resolution [Hansen et al., 2018], cellular modifications present in disease states [Li et al., 2016], the organization of the actin cytoskeleton in neuronal axons and dendrites [Xu, Zhong and Zhuang, 2013], the structure of receptors and scaffolding proteins at synapses [Specht et al., 2013], and the study of the protein complexes that form the nuclear pore [Szymborska et al., 2013], to cite a few examples. In this article, we focus on two of the key statistical challenges in super-resolution data analysis: the localization of individual point-source fluorophores and the estimation of the number of fluorophores present in the sample. These tasks must be carried out with only rough prior knowledge of the number of activated fluorophores and the dynamics of their activation.

Of the many techniques that have been developed to achieve super-resolution imaging, we focus on a class of methods that are generally referred to as *single-molecule localization microscopy* (SMLM). These techniques include stochastic optical reconstruction microscopy (STORM) [Rust, Bates and Zhuang, 2006], photoactivation localization microscopy (PALM) [Betzig et al., 2006] and other variants. SMLM techniques rely on the ability of photo-activatable fluorescent proteins [Lippincott-Schwartz and Patterson, 2009] or photo-switchable fluorophores [Heilemann et al., 2005; Dempsey et al., 2011; van de Linde, Heilemann and Sauer, 2012] to alternate between a fluorescence-emitting state and a dark state. By only activating a subset of the total number of fluorophores within the field of view, the emitting molecules can be individually localized, thereby breaking the diffraction limit of light.

In principle, SRM is poised to enable accurate counting of single molecules, thereby permitting the study of protein stoichiometry and dynamics under physiological conditions. Counting is, however, highly dependent on the characteristics of the molecule used as a fluorescent tag, and, in order to obtain an accurate count, several obstacles must be overcome. For example, fluorophore photophysics result in molecule over-counting when fluorescent molecules “blink” by transiently alternating between non-emitting dark and emitting light states [Heilemann et al., 2009; Roy et al., 2011]. Even more troublesome is the possibility that photoconvertible probes can be reactivated after a lengthy stay in the dark state [Annibale et al., 2010, 2011]. Furthermore, the actual number of photoactivatable probes, referred to as the *photoactivation efficiency*, is a missing parameter in all SMLM experiments that—in the absence of a second fluorescent probe that does not blink—can only be corrected computationally [Durisic et al., 2014; Zanacchi et al., 2017]. Many algorithms have been devised to compensate for blinking, ranging from semi-empirical approaches [Annibale et al., 2011] to more robust procedures that account for a single dark state [Lee et al., 2012; Rollins et al., 2015], many dark states [Hummer, Fricke and Heilemann, 2016] or the presence of many fluorophores and binding sites within a diffraction limited spot [Nino et al., 2017]. An ideal workflow designed to analyze SRM data sets would be capable of accounting for the different blinking properties of fluorophores while separating each fluorophore in space.

We devise a statistical approach to analyze STORM or PALM localizations computed by any conventional detection software [Holden, Uphoff and Kapanidis, 2011; Sergé et al., 2008; Ovesnỳ et al., 2014] and to infer the most likely fluorophore arrangement in the field of view. We model the observations as arising from a Poisson process endowed with a constant background density[Taddy et al., 2012; Kottas and Sansó, 2007] and we use Bayesian non-parametric (BNP) priors to model the number of fluorophores present in the sample [Blei et al., 2006; Broderick et al., 2013; Huggins and Wood, 2014]. Our approach involves two stages. First, we collapse observations across time and use a Dirichlet process (DP) mixture model to infer fluorophore events in the field of view. The DP is a BNP prior with the flexibility of accounting for an unbounded number of fluorophores. The model is scalable and fully amenable to approximate Bayesian posterior inference through the use of variational mean field methods. Moreover, by exploiting the locality of the problem and using a quadtree [Finkel and Bentley, 1974] data structure, we are able to build a high-performance implementation capable of analyzing millions of points in minutes on a standard laptop computer. Posterior inference using this model creates proposals for more complex algorithms.

Second, to model time dependencies between fluorophore observations, we incorporate the photo-physical properties of the fluorophores. We incorporate a hidden Markov model (HMM), which has been shown previously to accurately reflect fluorophore dynamics when the states of the fluorescent molecules are described by transitions between an inactivated state, a light-emitting state, a dark state resulting from radiating decay, and, finally, a bleach state in which the fluorescent molecule is unable to emit light [Lee et al., 2012]. To discover fluorophore positions and to assign observations to each active fluorophore jointly, we propose a model based on the Markovian Indian Buffet process [Gael, Teh and Ghahramani, 2009]. The entire set of fluorophores is shared among observations present at each time-point. Individually, however, only a subset of fluorophores is active and able to emit observations at each time-point. Our model decouples the probability of fluorophore appearance at each time point from the probability that an observation is assigned to the fluorophore. By doing this, we are able to assign many observations to each active fluorophore (presumably due to duplicate fluorophore localizations) and to account for background noise (i.e., spurious observations).

The paper is organized as follows. In Section 2, we begin by presenting the SMLM data sets analyzed in this work. In Section 3 we review the Dirichlet process and discuss a time-independent model for pointillistic data that has been pre-processed by deconvolution algorithms and we present a mean field inference algorithm to estimate the parameters of the model. Then, in Section 4, we extend the model to include fluorophore photophysics and create a time-dependent formulation based on the Markovian Indian Buffet process. In Section 5, we describe related work. In Section 6 we explore the performance of our algorithms on biological data sets and assess the accuracy of our method. We conclude in Section 8.

A list of notational conventions and further details on our algorithms and experiments can be found in the Supplementary Material [Gabitto et al., 2019].

## 2. Experimental Datasets

In this work we consider real biological data sets obtained using STORM imaging. The first data set consists of a 3D DNA origami scaffold equipped with multiple handles for the attachment of different molecules [Zanacchi et al., 2017]. The scaffold is 225 nm long and consists of a 12-helix bundle with 6 inner and 6 outer helices. It contains 15 attachment points, separated by a distance of 14 nm, that project outward and provide site- and sequence-specific positions to which fluorophores or proteins of interest can be functionalized (Figure 1A). First, at handle position 14, TAMRA fluorophores are attached to enable identification of the DNA scaffold under wide field imaging. Next, complementary handle sequences labeled with Alexa Fluor 647 were attached to handles 1, 7 and 13 of helix 0 to permit identification of single fluorophores (Figure 1B-D). STORM data sets generated using the scaffold were kindly provided by the authors of the original work [Zanacchi et al., 2017] for the demonstration of our method. We localized individual molecules present in single frames and used these positions as the input to our analysis. A second data set consist of super-resolution STORM imaging of the nuclear pore complex (NPC) in nuclear envelopes. NPCs provide access to the cell nucleus, thereby permitting the transport of proteins and RNA through the nuclear envelope. The function of the NPC is not limited to molecular trafficking, they are also involved in diverse cellular processes [D’Angelo and Hetzer, 2008]. The nuclear pore possesses a highly stereotyped configuration: proteins within NPCs are arranged in an eight-fold symmetric, cylindrical assembly consisting of approximately 30 different proteins of the nucleoporin (Nup) family [Kim et al., 2018]. Here, we analyzed a nuclear pore complex data set in which the Nup-107 protein is tagged with Alexa Fluor 647 in the nuclear pore membrane of U-2 OS cells and imaged using dSTORM on a commercial Leica SR GSD 3D microscope (kindly provided by the Reiss Lab, data from Li et al. [2018])(Figure 2A). Nup-107 proteins belong to the best-studied module within the NPC, the Y-complex (its name describes the shape in which proteins in the module assemble). Nup-107 surrounds the NPC, localizing to the nuclear rim and forming an eight-fold symmetrical ring [Beck and Hurt, 2017] (Figure 2B). This highly reproducible symmetrical ring provides a ground-truth structure to which we can compare in order to evaluate our algorithms.

**Fig 1.**
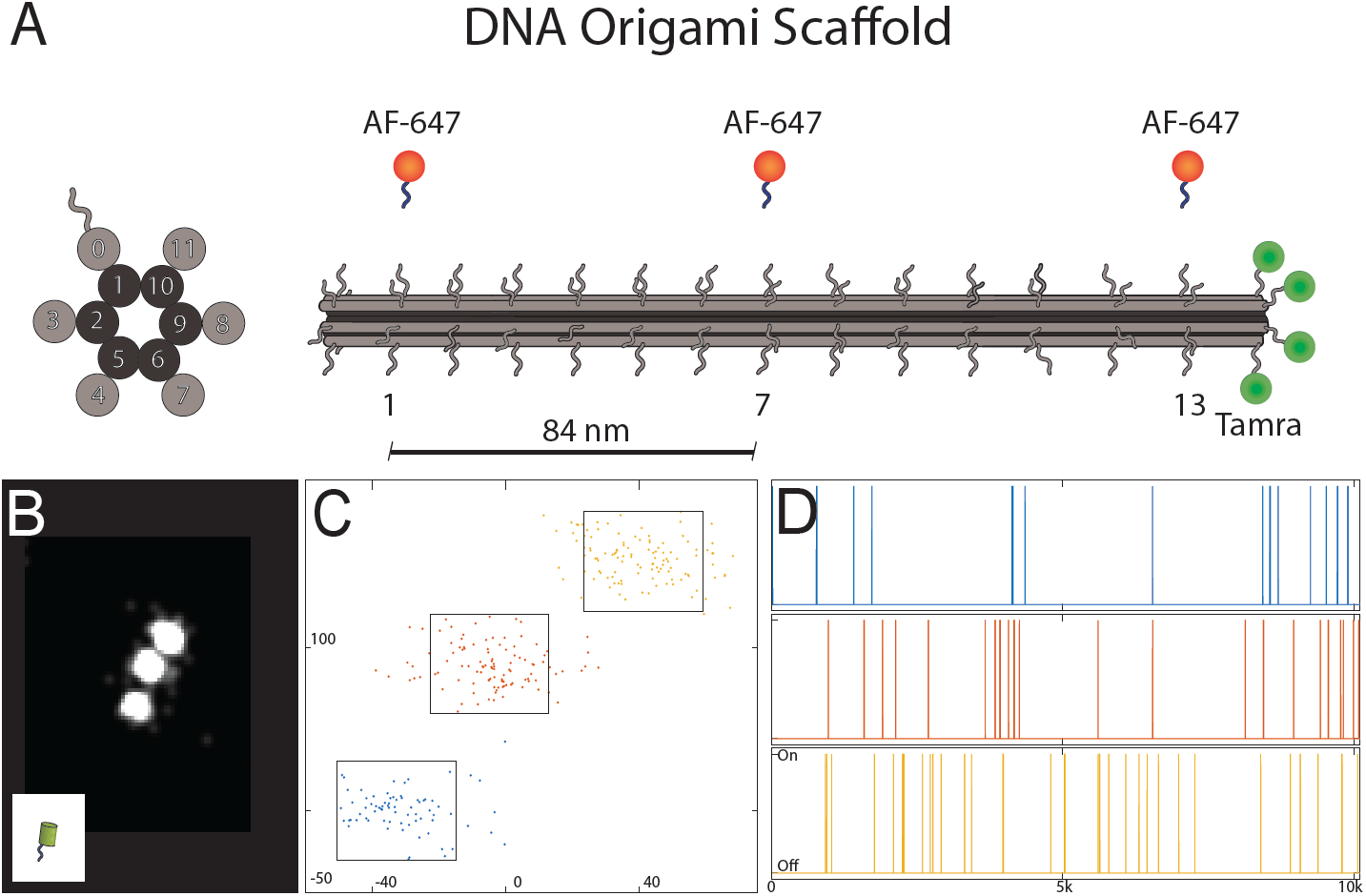
Super-resolution imaging of a DNA origami scaffold. a) DNA origami scaffold representation. Left, cross section depicting the double helix barrels, sequence specific handles protrude outward from outside helices. Right, the scaffold is 225 nm long and possess 15 handles. Fluorophores with handle complementary sequence are attached to handles 1, 7 and 13. At handle position 14, TAMRA fluorophores are joined for visualization under wide field fluorescent microscopes. b-c-d) Example of super resolution imaging of DNA origami scaffold in which sequence specific Alex Fluor 647 fluorophores are attached to handles 1,7 and 13. b) Fluorophore localizations smoothed with a Gaussian kernel (width = 10nm) and aggregated across the time series. c) x-y positions of fluorophore observations. d) Time traces depicting the frames in which each of the three fluorophores is observed.

**Fig 2.**
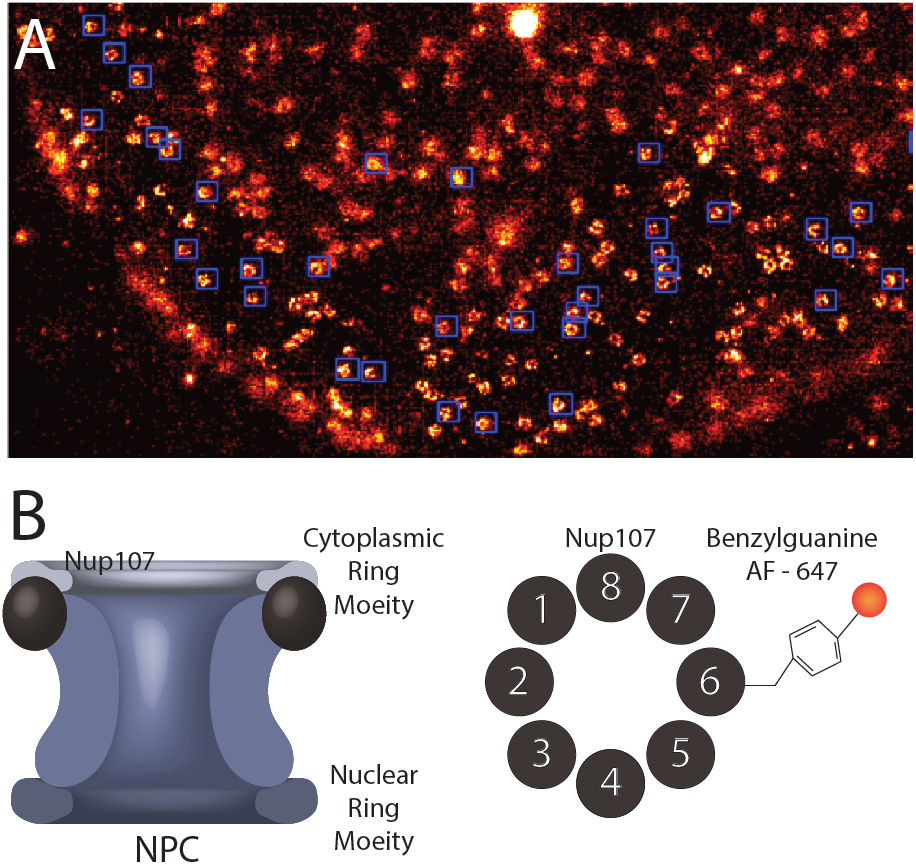
Super-resolution imaging of the nuclear pore complex. a) Example field of view in which nuclear pore complexes are imaged with super-resolution microscopy. Nuclear pore complexes are identified by template cross-correlation and enclosed by blue rectangles. b) Nuclear pore complex schematic. Left, cross-section of the nuclear pore complex on the nuclear membrane indicating in black, Nup-107. Right, top view of the nuclear pore complex highlighting Nup-107 octagonal symmetry labeled with Alexa Fluor 647.

## 3. A Time-Independent Model of Fluorophore Locations

A variety of existing techniques produce pointillist representations of fluorophore centers [Holden, Uphoff and Kapanidis, 2011; Sergé et al., 2008; Ovesnỳ et al., 2014]. These locations are prone to error due to their lack of correction for temporal effects (i.e., blinking) and their failure to integrate measurement uncertainty into the analysis. Moreover, spurious fluorophore locations can be created as the result of inaccurate modeling of the microscopic point spread function and the nuances of optimization algorithms. Our approach addresses these limitations within a Bayesian framework; moreover, to account for uncertainty in the number fluorescent proteins in the sample, our modeling formulation is based on a Bayesian nonparametric approach. We begin this section by briefly reviewing the BNP prior that we used to model fluorophore centers.

### 3.1. The Dirichlet Process

The Dirichlet process (DP) [Ferguson, 1973] can be understood as an infinite-dimensional analog of the Dirichlet distribution. The DP is a normalized random measure characterized by a scaling parameter *α* and a base measure *G*_0_ defined on the set Θ. We denote a draw from the DP as *G* ∼ DP(*α, G*_0_). The DP is defined by its possession of Dirichlet marginals; that is, for any finite partition *A*_1_, …, *A*_*K*_ of Θ, we have:

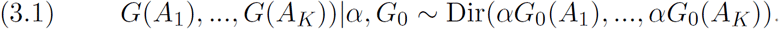

It can be shown that *G* is an atomic measure with probability one. Sethuraman [1994] provided an explicit representation of a draw from a DP, via a *stick-breaking construction*:

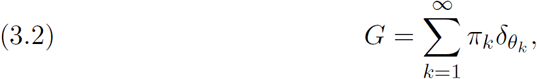

where 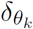 is an atom located at *θ*_*k*_ ∈ Θ, and the random weights *π*_*k*_ depend on *α* as follows:

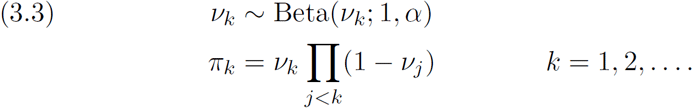

The random sequence (*π*_*k*_) sampled according to Equation (3.3) is said to follow a GEM distribution [Ewens, 1990].

The atomic nature of the DP has been exploited by many authors by using a DP as a prior for Bayesian mixture models. In this setting, the random weights of the atomic measure correspond to mixing proportions, and the locations of the atoms represent the parameters of the mixture components.

### 3.2. Model overview

Our observed data is a set of two- or three-dimensional fluorophore locations (i.e., putative fluorophores at different time frames). We begin our analysis by collapsing across time and inferring fluorophore centers in this collapsed data set. These inferences can be performed quickly, and they provide a seed for our more complex time-dependent model, described below in Section 4. Each observation *x*_*n*_ has an associated localization accuracy *σ*_*n*_ that informs confidence in the estimated location (*σ*_*n*_ is obtained by typical pre-processing steps). We model these observations as arising from a non-homogeneous spatial Poisson process defined on the observation box *R* = [*a*_1_, *b*_1_] × [*a*_2_, *b*_2_] × [*a*_3_, *b*_3_], with intensity *λ*(*x*) for *x* ∈ *R*. For such random processes, the following holds true:

i. For any bounded set *S* ∈ *R*, the number of points in *S* is Poisson distributed, 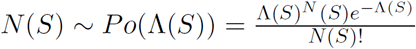, where Λ(*S*) = ∫ *λ*(*x*)*dx*.
ii. Given *N* (*S*), the point locations within *S* are i.i.d. with density 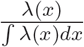.

The intensity *λ*(*x*) can be specified in terms of two independent factors— a total scalar intensity *λ*_0_ and a spatial density *f* (*x*), *λ*(*x*) = *λ*_0_*f* (*x*)— allowing the Poisson process likelihood to be written in separable form, thereby transforming the problem into one of density estimation:

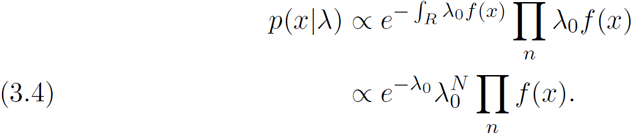

Placing a gamma prior on *λ*_0_ permits its inference in a manner that is independent of the spatial density. Next, we specify a prior for the process density *f* (*x*) with the domain restricted to the observation window *R*. We assume that every observation arises either from a constant background noise distribution or is a fluorophore which is distributed according to a truncated Gaussian distribution with mean *µ* denoting the fluorophore position. Building on Rubin-Delanchy et al. [2015], we propose a mixture of two components: spatially random background noise with constant density 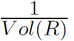, and a Dirichlet process containing an unbounded number of fluorophores. This prior is consistent with the idea that not every fluorophore is observed during the imaging experiment and that the number of fluorophores would increase if the experiment would have continued.

The graphical model representation is summarized in Figure 3. Defining 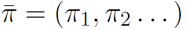, the overall model specification is summarized as follows:

**Fig 3.**
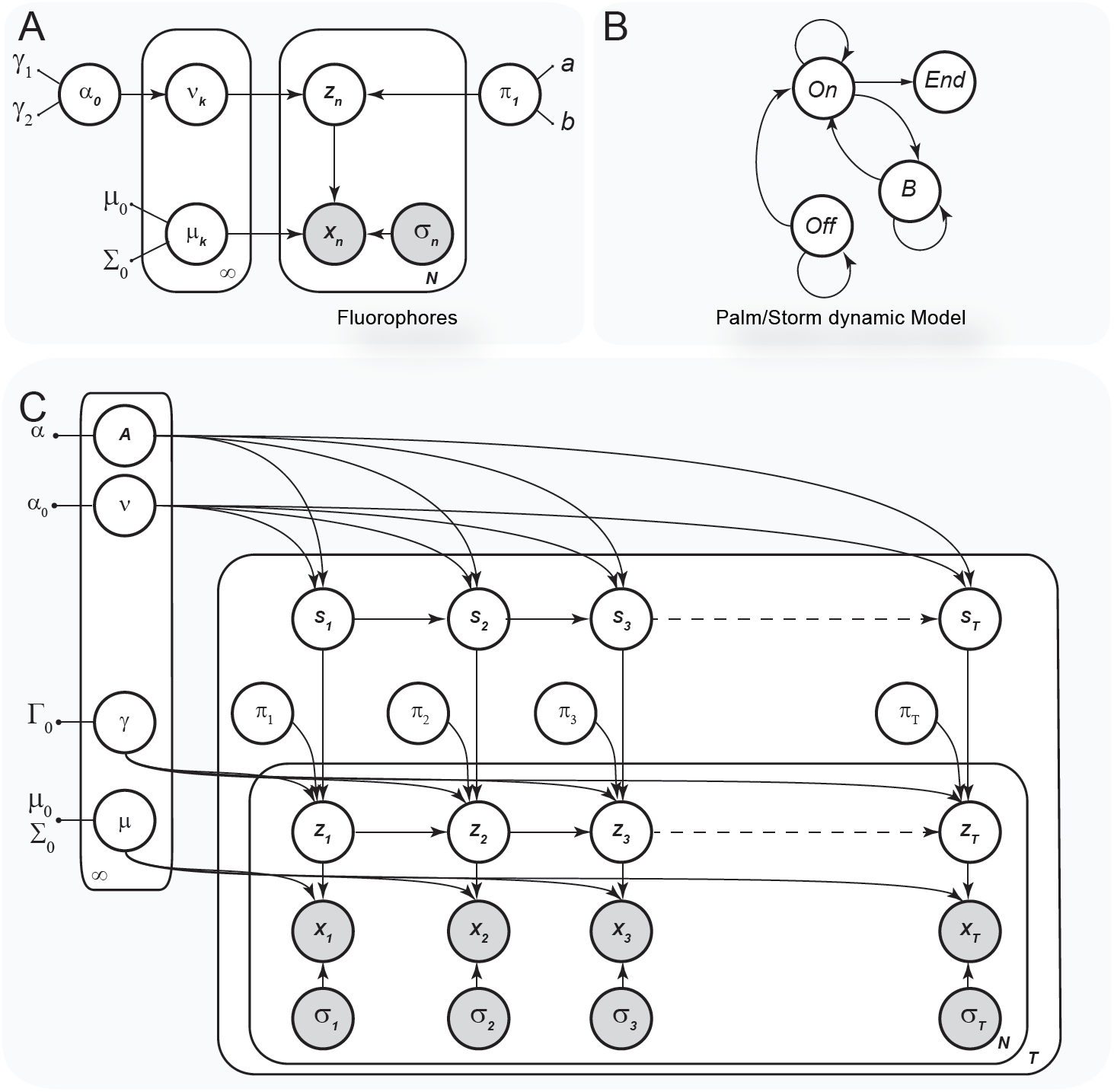
Time-independent and time-dependent Graphical Models for super-resolution localization. a)Time independent graphical model. b) Different states for the hidden Markov model used to model storm or palm data sets. c) Time dependent graphical model.

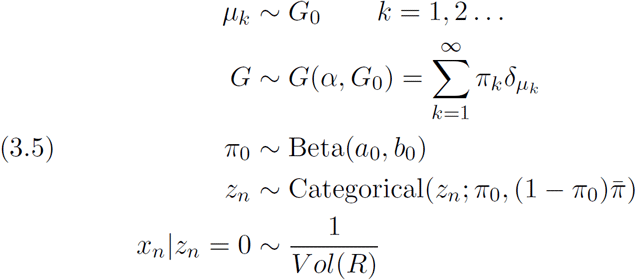

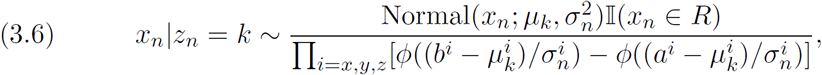

where *n* ∈ {1, …, *N*} in the last three equations, 𝕀 is an indicator function, *ϕ* is the standard Gaussian cumulative function needed to normalize the truncated Gaussian and 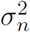 denotes a diagonal covariance matrix which ensures the factorization of the normalization constant. These modeling assumptions define a joint distribution *p*(*x, z, µ, ν*|*σ*) on the data and the latent structure of our model, where we use *ν* for the stick-breaking representation (3.3). After observing data (*x, σ*), our inferential goal is to obtain the posterior distribution of the fluorophore locations *µ* and assignments *z*. We refer to *µ* and the parameters of the stick-breaking prior as global parameters because they generalize to new observations, in contrast to the states *z*_*n*_, which are local to a specific observation.

### 3.3. Mean field variational inference

Exact posterior inference is intractable in our model. We therefore resort to computing an approximation to the posterior distribution using a variational mean field method [Wainwright et al., 2008]. Let *Q* be a family of distributions on Θ = (*π*_0_, *ν, z, µ*), the space of latent variables. For *q* ∈ *Q*, we search for a family of distributions that maximizes the evidence lower bound (“ELBO” = ℒ (*q*)):

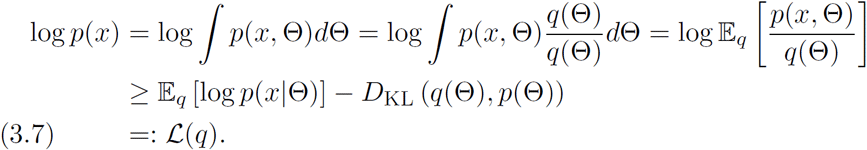

We seek a distribution *q* over the latent variables that is close to the true posterior and also lies within a factorized family, *q*(Θ) = *q*(*π*_0_)*q*(*ν*)*q*(*µ*)*q*(*z*). Each of the factors belongs to a particular member of the exponential family, except for the truncated normal distribution that characterizes the fluorophore centers. For the latter, we make use of the fact that our problem contains strong spatial information. In particular, we possess a priori information regarding the scale of the variance of each fluorophore center, given by the average uncertainty of the observations. By augmenting the size of the bounding box, we can assume the terms 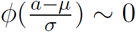 and 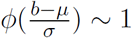, removing the need to explicitly truncate. Numerical exploration of the validity of this approximation is performed in the supplemental material [Gabitto et al., 2019].

By approximating the generative model in an unconstrained space, we restrict our subsequent analysis to a variational distribution with the following factorized density:

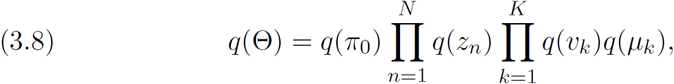

where

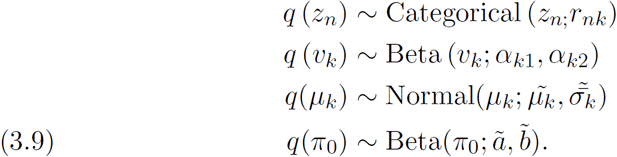

### 3.4. Inference

The computational task for our posterior inference algorithm is to find a set of parameters that maximizes the ELBO. Our algorithm updates free parameters in the variational distribution via coordinate ascent variational inference (CAVI). We present updates for global and local factors that converge to a local maximum. To simplify calculations, the ELBO is arranged into three terms that account for data generation, global variables characterizing our stick-breaking construction and an entropic term:

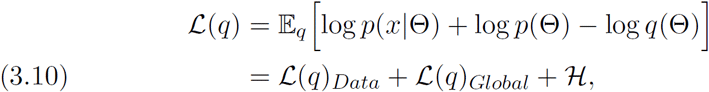

where

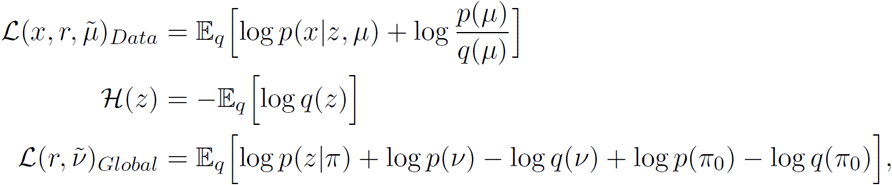

and where all expectations are taken according to the variational distribution. Due to the conjugate exponential family terms in the ELBO, the CAVI updates are easy to compute; see the supplemental material for the details [Gabitto et al., 2019].

Finally, we treat the hyper-parameter *α*_0_ as random in both the generative model and the variational distribution [Blei et al., 2006]:

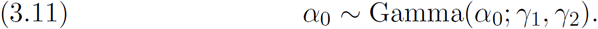

### 3.5. Scalable inference by exploiting spatial constraints

Standard variational inference assigns to each data point a positive posterior probability *r*_*nk*_, effectively assuming that each data can arise from any fluorophore. This instantiates matrices that demand dense memory storage and computation that scales with the total number of clusters. Our problem presents certain advantages due to the locality of its fluorophore assignments. This advantage translates into near certainty that only a few clusters have meaningful posterior mass for any given observation. Unlike approaches that instantiate a fixed number of clusters [Rubin-Delanchy et al., 2015], our approach assigns non-zero mass to clusters residing within a certain spatial distance of each observation; this distance acts as a threshold and is the only tunable parameter.

We harness the spatial locality to speed up computations by making use of a *quadtree*, a tree-like data structure that recursively subdivides two-dimensional space into four quadrants [Finkel and Bentley, 1974]. We exploit quadtree decomposition during the computation of local assignments. This step computes the posterior probability of assigning an observation *n* to fluorophore *k* for each observation *r*_*nk*_. Our CAVI algorithm optimizes 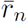 by fixing global fluorophore parameters according to the following objective:

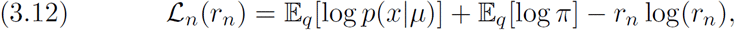

subject to the constraint that ∑_*k*_ *r*_*n,k*_ = 1 and *r*_*n,k*_ > 0. The first two terms represent the log posterior assignment of observation *n* to cluster *k*. Following Rubin-Delanchy et al. [2015], we replace the variational objective with a new one that dynamically limits the number of clusters to which an observation can be assigned. Instead of instantiating *r*_*n*_ as a dense vector, we compute only the non-zero mass entries for each observation n as determined by the clusters found within a certain distance of the observation. The new optimization problem can be written as:

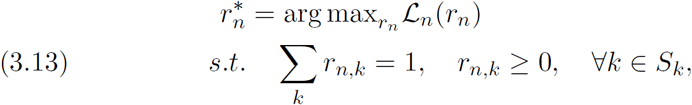

where the set *S*_*k*_ is computed dynamically at each iteration by querying a quadtree structure built on the locations of the *K* fluorophores. Observations that return no fluorophore centers are automatically designated as noise. Selecting the threshold distance results in a tradeoff between execution speed and inferential accuracy. In our case, the tradeoff is a favorable one, due to the strong locality of the observations arising from each cluster. This new constrained optimization problem can be solved by exponentiation of the log posterior assignment and normalization of the subset of active fluorophores. See Figure 4 for a numerical example of quadtree scaling.

**Fig 4.**
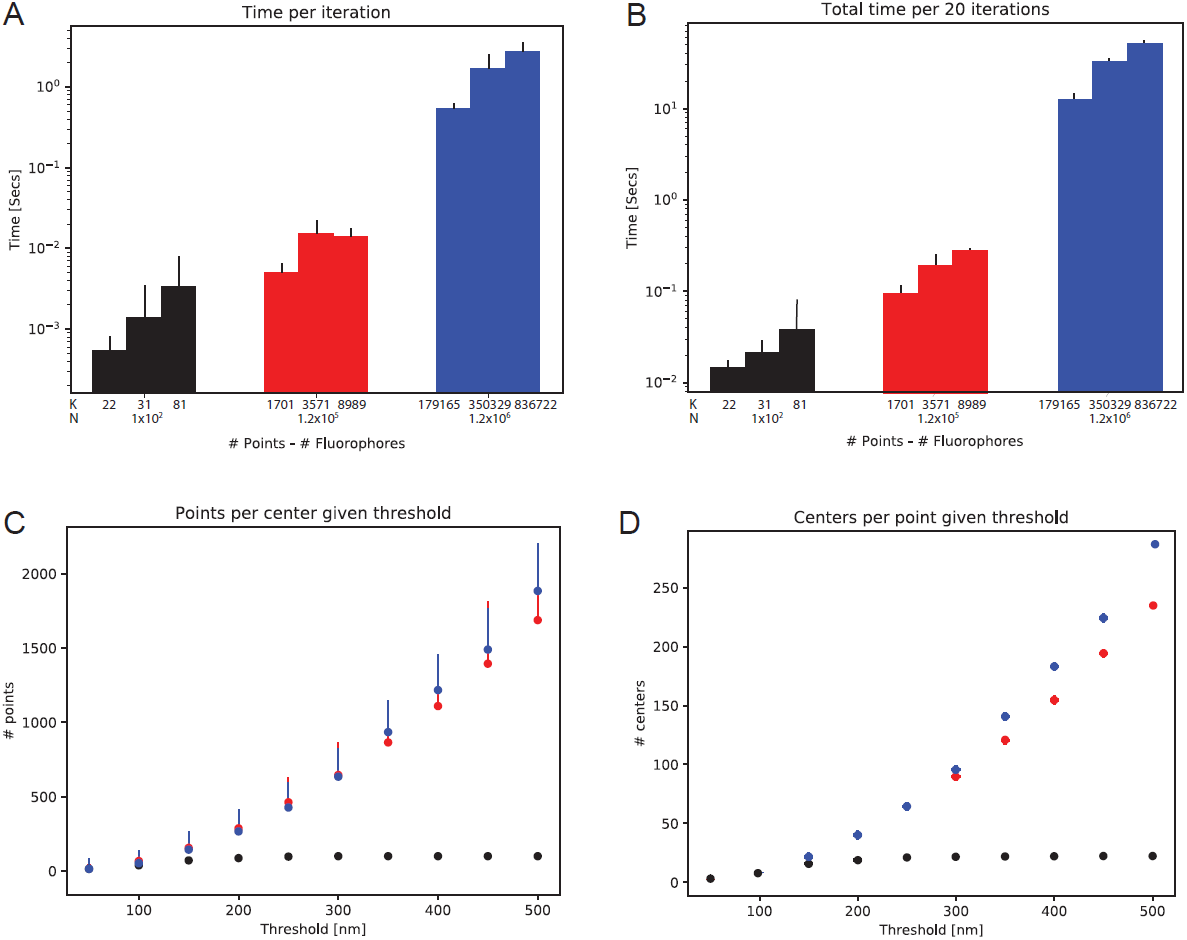
Performance of quadtree acceleration. For three different data sets, containing 100, 12 × 10^3^ and 1.2 × 10^6^ points, respectively, the algorithm was initialized with three initial conditions that gave rise to different numbers of initial clusters. a) Computational time for one iteration for each of the three data sets. b) Total time to compute 20 complete iterations, including local and global steps. By varying the threshold distance every quadtree query returns a different number of clusters k′ < K for each point. c) Average number of points per center and d) average number of centers per point. Thresholds lower than 150 nm return an equal number of centers independently of the size of the data set.

### 3.6. Reliable Bayesian inference via state space adaptation

Variational inference algorithms converge to local optima. We thus implement a multiple trial procedure that yields an algorithm that has the flexibility to improve initial assignment estimates. Specifically, we develop a series of fluorophore proposals aimed to obtain improvements in the ELBO. We interleave proposals that randomly split or merge observations assigned to fluorophores. We also create proposals that create or delete fluorophores by removing or assigning points to noise. To evaluate proposals rapidly, we simplify ELBO calculations for the gap between old and new fluorophores’ configurations. These calculations are reproduced in the supplementary material [Gabitto et al., 2019].

## 4. A time dependent model of fluorophore locations

We turn to the second phase of our single molecule localization procedure. Here we consider each individual observation in space and time, taking into account the photo-physical properties of each emitting molecule. In this formulation, a large number of fluorescent molecules is present in the sample but only a fraction of them are visible at each time point. We design a statistical model capable of analyzing spatio-temporal localization by relating observations at each time point to a collection of *K* fluorescent time series, where *K* is unknown and subject to posterior inference. Major challenges include the need to distinguish observations from background noise and the possibility of assigning more than one observation to a given time series at each time point.

To address these challenges we based our model on a Bayensian nonparametric prior representing an unbounded number of components, each consisting of a Markov chain. At each time point, observations can be assigned to only one of the active chains. Our model is based on the Markovian Indian Buffet process (M-IBP)[Gael, Teh and Ghahramani, 2009; Valera, Ruiz and Perez-Cruz, 2015] together with a time-dependent Dirichlet distribution.

### 4.1. A dynamic prior that shares features across time points

In this section we derive a prior distribution over binary matrices with a finite number of columns, each column representing a Markov chain model of a fluorophore. This formulation is closely related to the M-IBP, having a temporal dynamics that is specialized to the time evolution of fluorophores. We begin by introducing a latent state variable 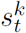, representing the state of the fluorophore *k* at time point *t*. The dynamics of this variable is governed by a transition matrix *A* such that:

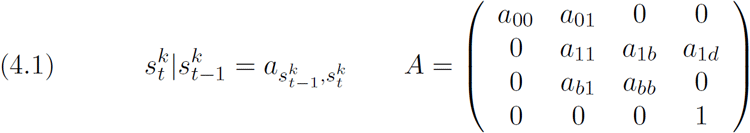

with prior distributions given by:

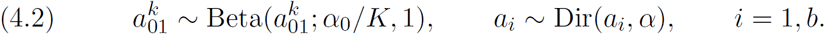

The transition matrix models an inactive state (0), a light-emitting state (1), a blinking state (b) and an dead state (d). Next, we introduce a feature vector 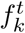 associated with the presence (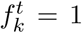 or 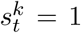, on) or absence (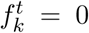 or 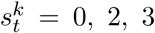 inactive, blinking or dead) of fluorophore *k* at time point *t*. This feature vector represents a binary matrix having as many columns as there are fluorophores present in the sample. Each feature vector follows a time evolution generated through the latent states:

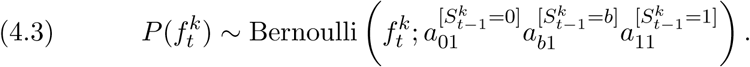

By introducing a set of count variables, *c*, to indicate the number of transitions between two states (e.g., *c*_01_ is the number of transitions from the state 0 to 1) we can write the probability of the entire binary matrix as follows:

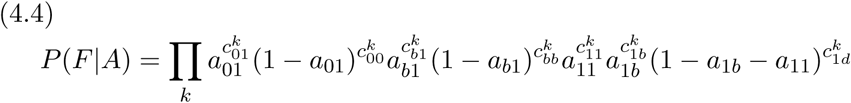

In a similar vein as the M-IBP, we calculate the marginal over the matrix *F* by integrating out transition probabilities:

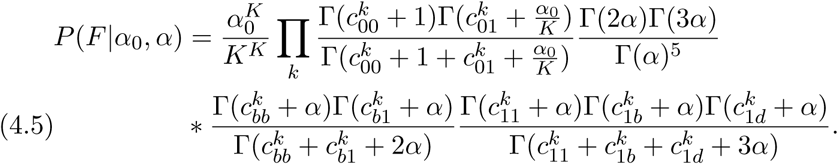

This represents the probability of each binary matrix defined by our time-dependent prior. Next, we use this matrix to assign observations at each time point to one of the active chains 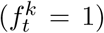. We do so by associating to each chain a gamma variable (*γ*^*k*^). Then, at each time point, we form a Dirichlet distribution by normalizing the gamma variables over the space of active chains:

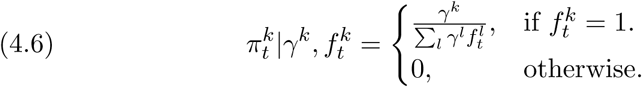

Finally, observations at time *t* are assigned to active chains by using a categorical distribution (*cat*(*π*_*t*_)).

In the supplemental material we show the infinite, i.e., nonparametric, limit of our model [Gabitto et al., 2019]. To obtain this model, we need to define two infinite mathematical constructs. In the first one, we derive a BNP prior over binary matrices that follows the fluorophore dynamics. The prior is exchangeable in the columns and it is also Markov exchangeable in the rows. In the second construct, we use the language of completely random measures (CRM) to show that a thinned CRM based on the Gamma process represents our model in the infinite case [Foti et al., 2013].

### 4.2. Model overview

Our model relies on the Markovian dynamics of our time-dependent binary matrix prior to generate a sparse set of active fluorophores at each time point, from which observations can be drawn We present our time-dependent fluorophore model as a graphical model in Figure 3, and we provide the full specification as follows:

1. Draw parameters:

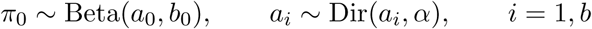
2. For *k* = 1 …*K*, draw chain parameters:

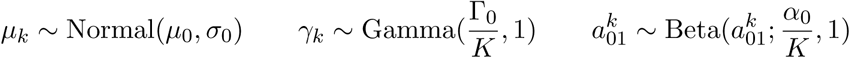
3. For each time point *t* = 1…*T*: For *k* = 1…*K*:

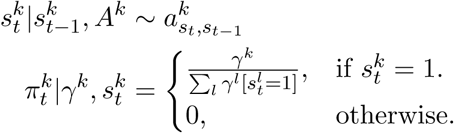 For *n*_*t*_ = 1…. *N*_*t*_:

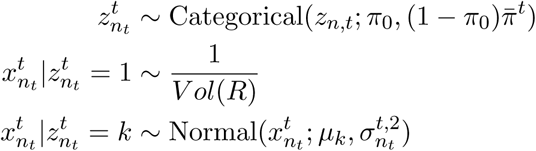

### 4.3. Mean field variational inference

In this section, we develop a variational inference algorithm to approximate the posterior distribution of the temporal model. We leverage the fact that a correspondence exists between the assignment of an observation at a particular time point to a fluorophore only if the fluorophore is active. Our approach employs the time-independent model as a seed, or initial condition, and we refine this solution through incremental move proposals. In particular, we propose a distribution from a factorized family, *q*(Θ) = *q*(*π*_0_)*q*(*µ*)*q*(*γ*)*q*(*z*)*q*(*S*)*q*(*a*), restricting our analysis to distributions *q* with the following dependence:

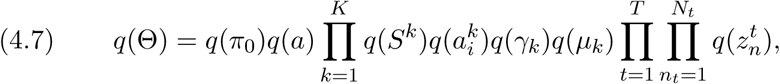

where

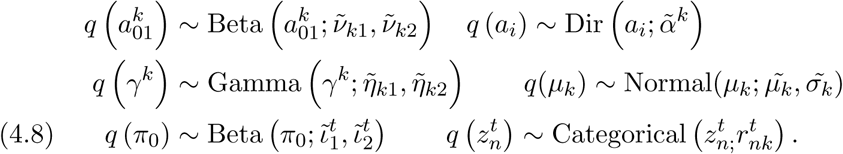

Lastly, we fit the state-variable dynamics via a structured variational proposal with Markovian structure [Hughes, Kim and Sudderth, 2015]:

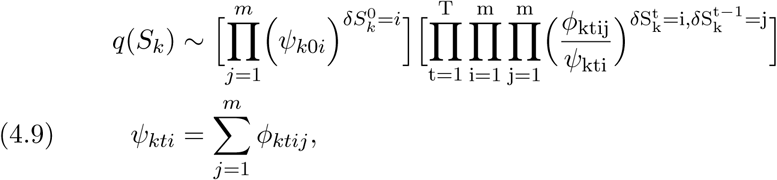

where the variational parameter *ϕ*_*ktij*_ represents the joint probability 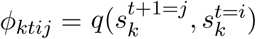 and *ψ*_*kti*_ defines the marginal probability 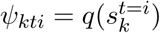.

### 4.4. Inference

We turn again to ELBO computation for the spatio-temporal model. In particular, we optimize the parameters of the fully factorized variational proposal amounts via coordinate ascent (CAVI). We arrange the ELBO into three terms accounting for data generation, the entropy and the KL divergences between our global parameter prior distributions and the corresponding variational proposals:

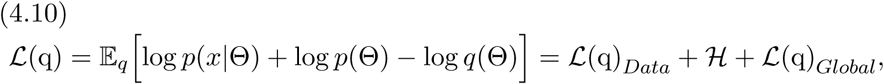

where

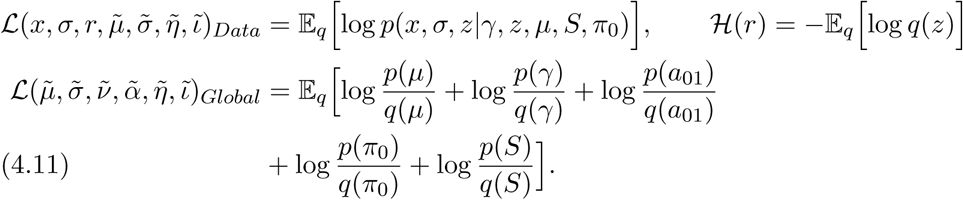

We compute the ELBO for this new model and take partial derivatives with respect to each variational parameter to derive the coordinate ascent updates. Most of the updates are straightforward to compute due to the conjugate exponential family factors. To optimize the local variables involved in the likelihood equation, we relax our generative model by introducing a variable *ϵ* ≪ 1:

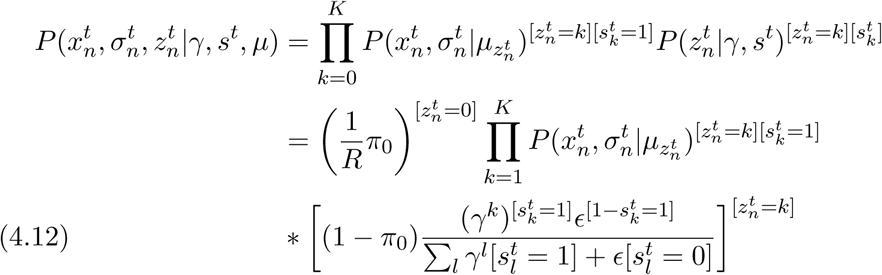

This relaxation smooths the ELBO and helps convergence. Finally, as noted by [Sun, Paisley and Liu, 2017] in a similar setting, the gamma normalization term presents computational difficulties, and we follow those authors in introducing an auxiliary variable *ξ*_*t*_ to further lower bound this term: [Sun, Paisley and Liu, 2017].

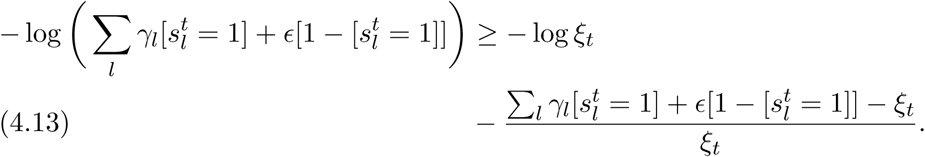

To obtain the value of *ξ*_*t*_, we differentiate the lower bound and set to zero, which yields the following update rule:

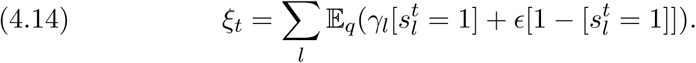

We reproduce the entire list of CAVI updates in the supplemental material [Gabitto et al., 2019].

### 4.5. Algorithmic work flow

To summarize, the analysis of a particular data set, based on a given set of fluorophore localizations in space and time, proceeds as follows. First, we collapse observation in time and select an initialization. We refine this initialization through our time-independent model. We interleave birth-death, split-merge proposals to explore different configurations of the state space and prune existing clusters. The final configuration of our time-independent algorithm seeds our time-dependent model. Finally, clusters are refined by split-merge moves that help to separate time-overlapping clusters. We approximate the entire time evolution of each fluorophore state when splitting or merging fluorophores in this model. To efficiently propose split moves, we calculate the posterior number of fluorophores inside clusters with an unusually high number of observations, and evaluate split proposals. The calculation of the posterior number of fluorophores given blinking statistics is detailed in the supplementary material [Gabitto et al., 2019]. Finally, at the end, for each fluorophore we compute the time evolution of each fluorescent trace.

## 5. Related work

There has been a great deal of previous work on the development to software to deconvolve super-resolution movies [Small and Stahlheber, 2014; Holden, Uphoff and Kapanidis, 2011; Sergé et al., 2008; Ovesnỳ et al., 2014]. Most of this work focuses on computationally efficient algorithms for detecting the Gaussian shape of the point spread functions without attempting to explicitly model latent temporal dynamics. Software that incorporates temporal information is extremely computationally costly and is prone to producing artifacts [Rosten, Jones and Cox, 2013]. Recently, deep generative models have been used to identify fluorophores from SR images, taking into account different PSF shapes [Sun, Archer and Paninski, 2017; Speiser, Turaga and Macke, 2019; Nehme et al., 2019]. Some of these approaches are complementary to ours and of similar computational complexity; others are significant more costly computationally. Our software builds upon fast single-frame deconvolution algorithms to incorporate temporal information into the localization analysis. Furthermore, we use spatially sensitive data structures to speed up calculations and facilitate scalability.

More broadly, identifying the number of different time series and assigning observations at each time point is a difficult task when the number of observations does not match the number of time series. This problem has been partially addressed by different authors. Several BNP approaches that capture time evolution—most based on the Hierarchical Dirichlet Process (HDP) [Teh et al., 2006]—have been developed in the setting of topic models. The HDP assumes that the probability of topics and the proportion of words explained within each document are coupled. This is undesirable, however, when there are rare topics explained by a large proportion of words in a small number of documents. A similar problem has been encountered in sparse topic modeling [Williamson et al., 2010; Faisal et al., 2012; Archambeau, Lakshminarayanan and Bouchard, 2014] where it is important to distinguish the probability that a topic belongs to a document from the probability of inclusion of the topic into the analysis. To address this issue, the authors proposed the use of a compound Indian buffet process-Dirichlet process. However, their work did not consider the dependency of the features across time and is limited by the sampling scheme developed.

The use of Bayesian nonparametric feature models for the modeling of time series was initiated by Fox et al. [2009]. Their work focused on motion capture data, a domain without the complexities and the scale of single-molecule imaging domain that is our focus. There has been significant follow-up work in this vein, including the use of a time-dependent beta process as a Bayesian nonparametric prior for feature allocation models [Perrone et al., 2016]. Again, however, the focus has been on small-scale problems and the methods are not directly applicable to the single-molecule imaging problem.

## 6. Realistic Simulation Studies

In this section we present studies assessing the performance of our algorithm on data sets derived from the DNA origami platform and the nuclear pore complex. Both data sets were presented in Section 2.

### 6.1. Data preprocessing and data sets construction

To create realistic SMLM data sets we make use of the DNA origami platform introduced in Section 2. Raw images of every data set were preprocessed with Thunder-storm [Ovesnỳ et al., 2014]. Briefly, raw images were imported into FIJI [Schindelin et al., 2012] and the Thunderstorm plugin was run with camera parameters and default approximate and sub-pixel molecule localization parameters. Next, observations with an unusual variance, uncertainty or intensity value (five standard deviations above or below the mean) were filtered out. We use these raw localizations to generate realistic data sets.

We isolated single fluorophores present in the data set in which fluorophores are attached to handle complementary sequences (Figure 2A). We verified that observations are localized with the reference TAMRA signal, that three sets of cloud points co-localized and that cloud distances were close to 84 nm (distance between handles 1-7 and 7-13). When these conditions were met, we then isolated individual cloud of points and considered each of them as an isolated fluorophore. For each extracted fluorophore, we computed its posterior fluorophore location according to our algorithm for just one fluorophore. This is the ground-truth fluorophore location against which we test our algorithm.

### 6.2. Model hyperparameters

For all subsequent computational experiments, we used weakly informative hyperpriors. We placed a Gamma(1, 0.01) prior on the concentration parameters *α*_0_. The parameters of the base measure were set from the data, with *µ*_0_ was chosen to be the center of the field of view and *σ*_0_ the maximum distance between observations. To set the prior on *π*_0_ we reasoned that in a real experiment, most of the points arise from real fluorophores. Accordingly we chose *a*_0_ = 1, *b*_0_ = 100. We chose the augmentation factor of the bounding box to be 1.25 times the average standard deviation of the points. This value seemed to be robust across the different data sets that we analyzed. For the time-dependent model, G_0_ was given a Gamma(100, 1) prior. The prior on the transition matrix was given via pseudo-counts *α*_*on,on/blink/dead*_ of 10, 5, 1 and *α*_*blink,on/blink*_ of 10, 1.

### 6.3. Computational experiments

In this section we present several different experimental scenarios that we employed to test the limits of our algorithm. In particular we aimed to assess 1) when our algorithm fails to distinguish nearby fluorophores; 2) how well our move proposals explore the ELBO, finding global optima; and 3) how well our procedures scales with an increasing number of fluorophores in the field of view.

To achieve this goal, we simulated fluorophore observations by randomly sampling a DNA origami fluorophore and placing its observations in a ground-truth position. To contaminate data sets with noise, we randomly selected observations from any fluorophore and randomly positioned them in the field of view. We quantified the noise level by measuring the ratio of noise points over points that belong a ground-truth fluorophore. By these means, we were able to construct realistic simulated data sets with realistic ground-truth observations. We judged performance with reference to a variety of metrics: the algorithm’s ability to correctly segment the data according to the underlying fluorophore location, the robustness to the algorithm to different number of fluorophores and noise level, and fluorophore detection based on fluorophore proximity.

### 6.4. Identifying nearby fluorophores

We simulated two fluorophores at different distances and different noise levels, and compared our algorithms (both the time-dependent and time-independent algorithms) to DBScan [Ester et al., 1996] Figure 6a. We explored the ELBO through different move proposals and returned the best configuration seen. In all cases, automatic DBScan settings explored through optics [Schubert et al., 2017] failed to identify the correct number of fluorophores. As seen in the figure, our algorithm performed significantly better than DBScan. Our algorithm performed poorly when observations are nearer than two times the observation standard deviation, although the dispersion of the localizations of each fluorophore also played a role in the performance. This limit is extended through the use of our time-dependent formulation. This limit is an intrinsic property of our model, as revealed by the average ELBO gap between the true fluorophore configuration and one in which fluorophores are merged Figure 6b. The gap trace switches sign at the limit, indicating a preferred incorrect fluorophore configuration. Our algorithm has a good performance above this limit that is robust to noise. Below this limit, even if data driven proposals can identified more than one fluorophore in the cloud of points, their location cannot be correctly determined.

**Fig 5.**
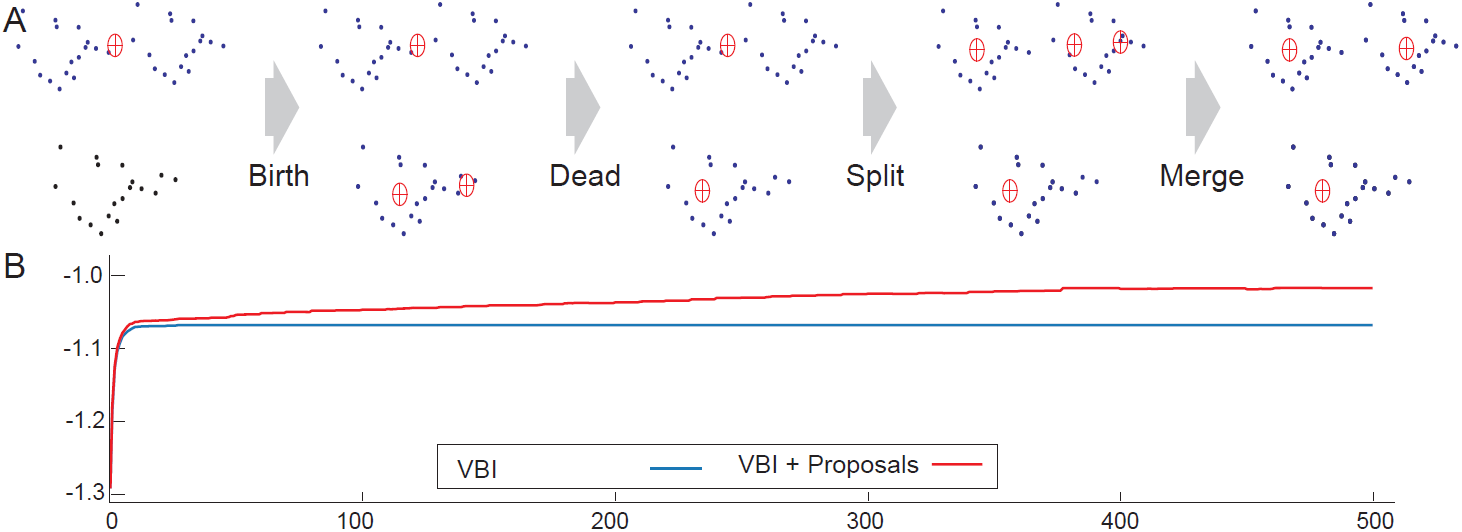
State space exploration through different proposal configurations. a) Cartoon representation showing how different configurations can evolve under birth, death, split, and merge proposals. b). ELBO evolution with and without state space exploration on a data set of 12000 points.

**Fig 6.**
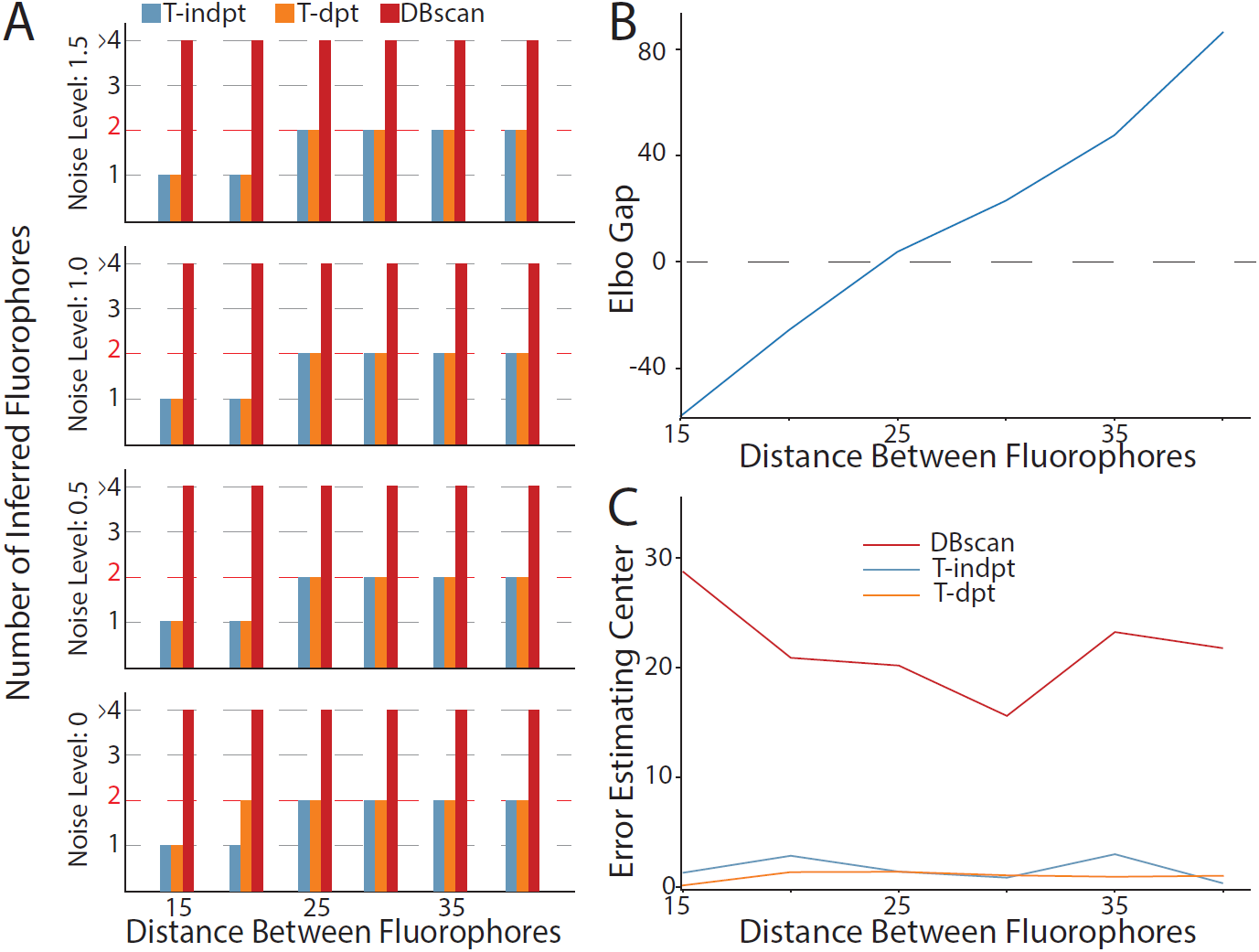
Realistic simulated experiments, distinguishing two individual fluorophores. a) Number of inferred fluorophores under different noise regimens, varying the distance between them. b) For a noise level of 0.6, ELBO gap computed as the difference between a model with 2 fluorophores and a model with 1 fluorophore. When the Gap becomes negative, the model prefers the incorrect configuration of just one fluorophore. c) Error of fluorophore inferred position.

We explored robustness to different noise scenarios, revealing that performance is maintained even when the noise level reaches a value of two. This noise corruption means that locally, two out of three points belonged to noise. Experimentally, it is highly unlikely to encounter such scenarios, and we did not observe it in our nuclear pore experiments. Finally, our time-dependent algorithm seems to be more accurate than our time-independent model in correctly localizing fluorophores’ true positions Figure 6c.

### 6.5. Scaling fluorophore numbers

Next, we simulated an increasing number of fluorophores at different distances, localizing them on a grid. We imposed a realistic level of noise of 0.5 Figure 7a-b. We varied fluorophore distances while remaining above our identification limit. We then applied our algorithms to every condition, randomly simulating the conditions six times and computing averages of inferred fluorophore numbers. As seen in Figure 7c, our algorithm is robust to an increasing number of fluorophores as long as we increase the number of proposals explored. There was a mild decrease in fluorophore recovery when the distances approaced the detection limit. As expected, true fluorophore localization degraded as fluorophores approached each other (see Figure 7d).

**Fig 7.**
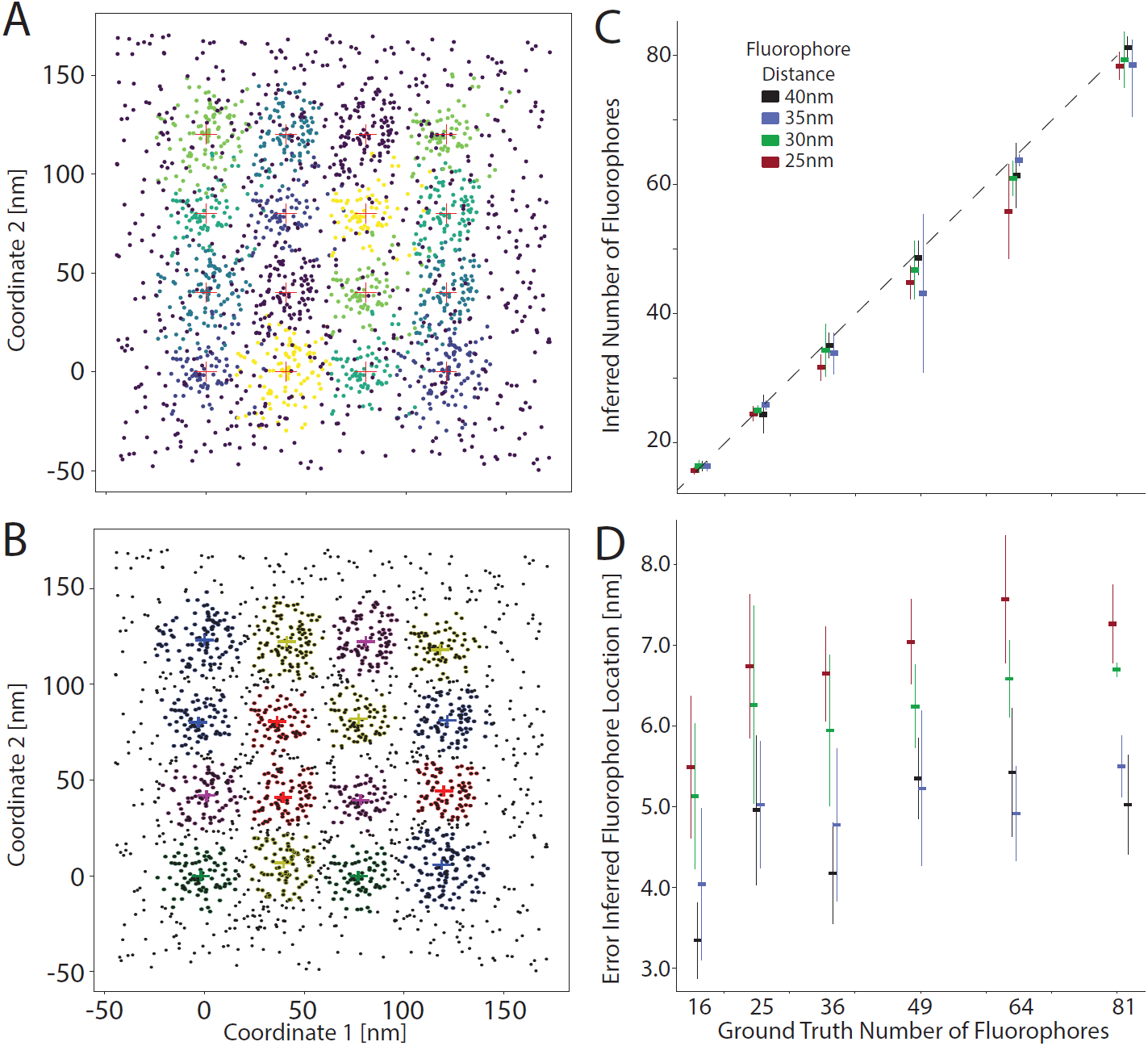
Realistic simulated experiments, scaling the number of fluorophores. a) True configuration for an example of 16 fluorophores in the field of view. b) For a configuration such as in a), inferred fluorophore positions. c) Number of inferred fluorophores when the ground truth number in the field of view is increased. d) Average error ± standard deviation for the inferred fluorophores’ positions.

## 7. Application to Nuclear Pore Complex Data

Finally, we present results of applying our algorithms to the nuclear pore complex, a real biological data set of known structure. To focus on the NPC, a field of view needs to be preprocessed, and NPCs isolated. To select several instances of the imaged NPCs, we proceeded by isolating a few candidates in the image and creating a searching template (Figure 8A). This template was cross-correlated against the entire image and candidates data sets were created from regions in which the correlation score exceeded a threshold (Figure 2A). These NPC datasets with localized emitters served as input to our analysis (examples of NPCs are reproduced in (Figure 8B)).

**Fig 8.**
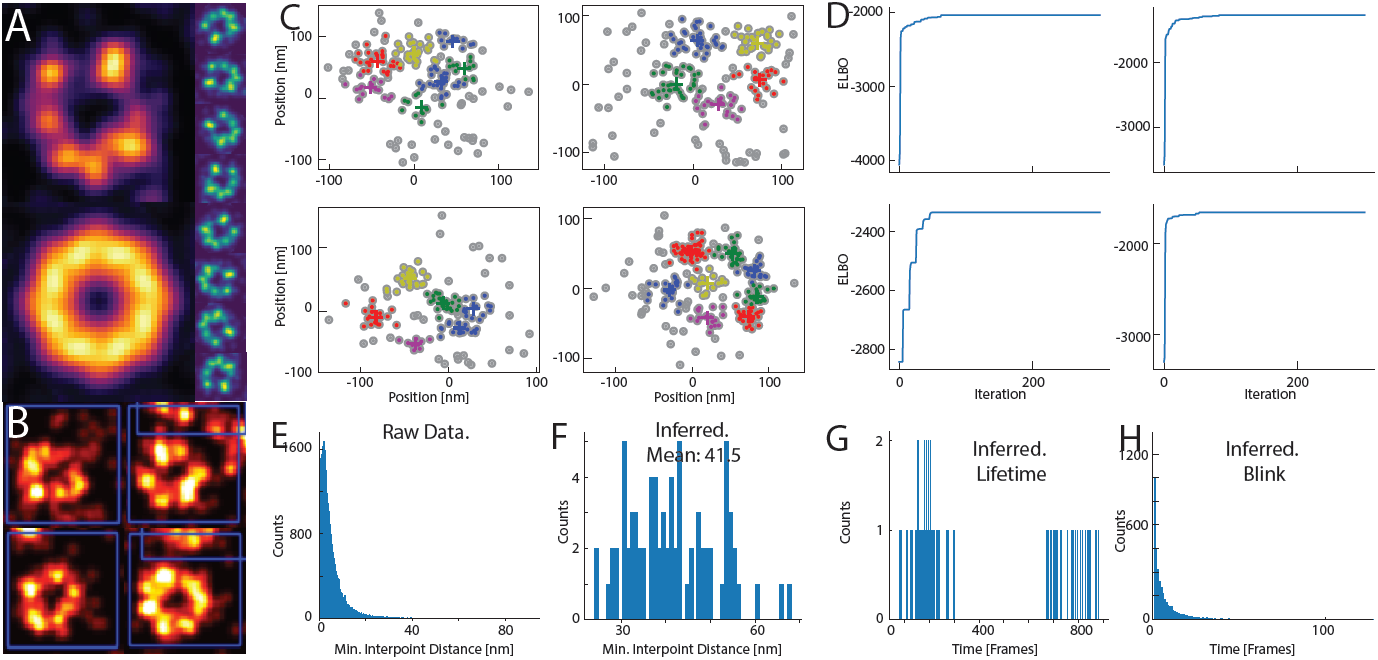
Analysis of Nuclear Pore Complex Datasets. a) Top left, a nuclear pore complex image convolved with a Gaussian filter. Bottom left, the nuclear pore complex is rotated eight times and these rotated images are aggregated to generate a nuclear pore complex template. Right, an eight-fold rotation of the nuclear pore complex. b) Nuclear pore complexes isolated through template matching. c) Localizations extracted from nuclear pore complex data sets. Inferred fluorophore centers highlighted with a cross in different colors. d) ELBO evolution during our inference procedure for the four datasets shown in c). e) A histogram depicting the distance for each point to its closest neighbor. f) Same as in c) but this time for each inferred fluorophore location. g) Inferred fluorophore durations. h) Inferred time interval between subsequent observations of each fluorophore.

We applied our algorithms to 369 NPC datasets, running them for a fixed budget of iterations (although convergence was recognized well before exhausting the budget) and identifying underlying fluorophores (Figure 8C-D)). We were specifically interested in assessing the validity of our inferred fluorophores. Therefore, we made use of the aforementioned NPC symmetry and counted fluorophore distance to the closest neighbor. Given the NPC octagonal shape, we thereby obtained a minimal fluorophore distance of 41.5nm, where the raw data minimal distance is close to 3nm (as illustrated in Figure 8e-f)). Moreover, as seen in Figure 8g-h, the inferred time dependent parameters were in accordance with published results, with the blinking probability exhibiting a clear exponential decay [Lee et al., 2012].

## 8. Discussion

We have presented a Bayesian nonparametric method for the identification of fluorescent molecules in super-resolution experiments. To obtain a procedure that is viable at the scale of realistic experiements, we developed a statistical methodology that proceeds in two phases. The first phase is based on a model that analyzes localization microscopy observations by collapsing temporal information. This model relies on the Dirichlet process as a prior on the underlying number of fluorophores present in the sample. To speed calculation, we used spatial data structures (quadtrees) to obtain individual fluorophore assignments. Inference in this model is performed using mean field variational inference, and the feature space is explored using state space adaptation techniques. Next, we developed a statistical model that incorporates temporal information into the analysis and accounts for fluorophore photo-physics. Taking the infinite limit of this model defines a nonparametric prior that is comprised of an infinite factorial hidden Markov model and a dependent Gamma process. This prior allows the assignment of different probabilities to the inclusion of a fluorophore at each time point and determines the probability of assigning an observation to each active fluorophore. To refine the inferred fluorophore numbers, we incorporate data-driven split-merge moves that split fluorophores based on fluorophore blinking statistics.

We demonstrated the utility of our model using realistic simulated data and a real data set in which the underlying biological structure is known. In this real data set, we were able to correctly infer fluorophore localization consistent with the geometry of the sample, as demonstrated by the inter-fluorophore position distributions. By using realistic simulated data, we showed that our model is robust to noise conditions encountered in real experiments. We expect our method to perform poorly in cases where fluorescent molecules are out of focus, which results in an inferred position that differs greatly from the true location.

A key feature of our algorithm is that it can be used as a postprocessor for any software pipeline that extracts raw fluorophore localizations from data. Our approach aims to integrate temporal information into the analysis and correct mistakes produced during single-molecule identification. We illustrated our method using the external software package Thunderstorm to process our images. In practice, there is evidence that such localization software produces mistakes when two nearby fluorescent molecules are active in the same frame due to a failure to correctly infere molecule locations. Future work could consider alternative ways of preprocessing imaging datasets. In this case, to further improve localization accuracy, it might be desirable to incorporate modeling of the point spread function into the model and to directly process raw images within a more sophisticated statistical framework.

Our method is implemented in python with a C++ kernel, available on github. Instructions on how to run the code as well as code to reproduce the results are available within the repository.

## Acknowledgments

We thank Melike Lakadamyali, Francesca Cella, Jonas Reis and Yiming Li for providing published datasets to validate our methods.

## SUPPLEMENTARY MATERIAL

**Supplement to “A Bayesian nonparametric approach to super-resolution single molecule localization”:**

(DOI: http://www.e-publications.org/ims/support/dowload/imsart-ims.zip). We provide additional material to support the results in this paper. We include in the supplemental materials, detailed derivations of parameter update rules, cluster refinement procedures and possible extensions of our algorithm to different time distributions.

